# Redefining catecholaminergic polymorphic ventricular tachycardia (CPVT) as a neurocardiac condition

**DOI:** 10.1101/2025.01.27.635037

**Authors:** Molly O’Reilly, Arie Verkerk, Carol Ann Remme

## Abstract

Catecholaminergic polymorphic ventricular tachycardia (CPVT) is an inherited arrhythmia syndrome characterised by adrenergic activity-induced sudden cardiac death. It is most often caused by mutations in the *RYR2* gene encoding ryanodine receptor 2 (RyR2), which is essential for intracellular calcium handling. Research has traditionally focused on the consequences of mutations at the cardiomyocyte level. However, RyR2 is also expressed in neuronal tissue, and patients often present with clinical signs of autonomic dysfunction. Here, we assessed if there is a neuronal phenotype in this classically cardiac condition.

Using the established CPVT mouse model *Ryr2*-R2474S, we found that RyR2 is abundantly expressed in stellate ganglia neurons (SGNs) - the adrenergic neurons that project directly to the myocardium and modulate heart function. We revealed that in isolated *Ryr2*-R2474S SGNs there is altered calcium homeostasis suggestive of intracellular calcium leakage, and increased neurite projections when maintained in culture. We furthermore showed that hearts of *Ryr2*-R2474S mice are sympathetically hyper-innervated, with an increased heterogeneity in innervation within the myocardium. This led to changes in the quantities of neurotransmitters and their metabolites within the heart, indicating increased norepinephrine turnover.

CPVT may therefore be redefined as a neurocardiac disorder, with neuronal dysregulation being a prominent component of the disease. In addition to cardiomyocytes, *RYR2* mutations also affect stellate ganglia neuron function and cardiac innervation, with potential implications for arrhythmogenesis, risk stratification and therapy.

**Graphical abstract:** 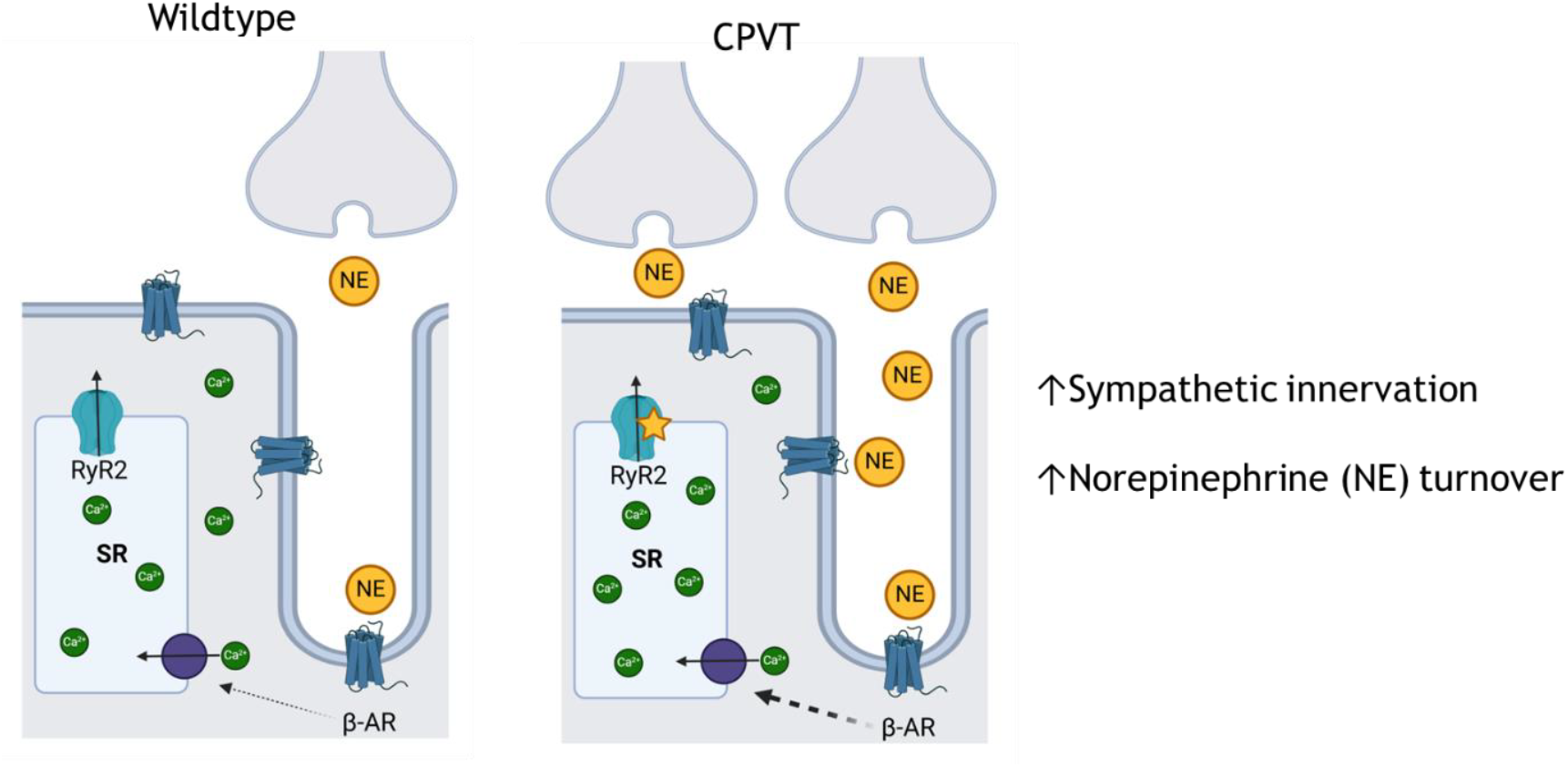

In wildtype hearts, activation of sympathetic neurons leads to their release of norepinephrine (NE), which binds to beta adrenergic receptors (β-ARs) on the cardiomyocyte membrane. The resultant initiation of intracellular signalling cascades involving cyclic AMP (cAMP) leads to reduced inhibition of the sarcoendoplasmic reticulum (SR) calcium ATPase (SERCA) pump and consequent calcium overload in the SR. This triggers the opening of ryanodine receptor 2 (RyR2), causing SR calcium to be released into the cytoplasmic space resulting in a calcium wave which triggers contraction.

In CPVT hearts, there is increased sympathetic neuron innervation of the ventricular myocardium. When cardiac sympathetic nerves are activated, there is an enhanced release of NE. This excess NE causes increased activation of β-ARs and an augmented intracellular cAMP response, which in turn leads to a greater reduction of the inhibition of SERCA, augmented SR calcium overload, and a heightened propensity for spontaneous calcium leakage to trigger a calcium wave, delayed afterdepolarisations, and arrhythmias.

Figure created with BioRender.com.

## Introduction

Catecholaminergic polymorphic ventricular tachycardia (CPVT) is an inherited arrhythmia syndrome that is characterised by adrenergic activity (exercise or emotion)-induced ventricular arrhythmias and sudden cardiac death (SCD) in young and otherwise healthy patients. It is a rare disease (1:5000/10000) but one that is extremely devastating – if left untreated, CPVT has a very high mortality rate, reaching 35% by 30 years of age (Kontula et al., 2005).

CPVT is most often caused by mutations in the *RYR2* gene encoding the ryanodine receptor 2 (RyR2), which is essential for intracellular calcium handling. In the heart, activation of sympathetic neurons leads to their release of the neurotransmitter norepinephrine (NE). NE binds to beta adrenergic receptors (β-ARs) on the cardiomyocyte membrane, which initiates complex intracellular signalling cascades involving adenylyl cyclase stimulation, cyclic AMP (cAMP) generation, protein kinase A activation and phospholamban (PLB) phosphorylation. The phosphorylation of PLB prevents it from inhibiting the sarcoplasmic reticulum (SR) calcium ATPase (SERCA) pump, which is responsible for transporting calcium into the SR. The reduced SERCA inhibition increases the rate of SR calcium uptake, leading to SR calcium overload, which is the trigger for RyR2 activation. Once a threshold level of SR calcium is reached, RyR2 transports calcium into the cytoplasmic space, causing a calcium wave that is responsible for cardiac contraction.

CPVT-related *RYR2* mutations are thought to increase the propensity for arrhythmias by modifying various aspects of this intracellular calcium signalling cascade (see Priori et al., 2021 for full review). A unifying theory has been proposed for all CPVT mutations (encompassing not only *RYR2* but also the other frequently associated gene, *CASQ2*). This proposes that SR store overload-induced calcium release (SOICR) is the common triggering mechanism, with the free SR luminal calcium level repeatedly exceeding the threshold for SOICR (MacLennan and Chen, 2009), leading to enhanced spontaneous calcium release which then triggers pro-arrhythmic calcium waves. Despite this knowledge, current therapy options are often not sufficient, and identification of high-risk patients remains challenging. This indicates that there are other mechanisms involved in the disease manifestation.

Research has traditionally focused on the functional consequences of mutations at the cardiomyocyte level. However, RyR2 is also expressed in neuronal tissue, being abundantly expressed in the brain (Abu-Omar et al., 2018). Affected patients typically present with clinical signs of autonomic nervous system (ANS) dysfunction, including bradycardia, syncope, disrupted heart rate variability and increased sensitivity to neuronal activity (Leenhardt et al., 1995; Postma et al., 2005; Azevedo et al., 2015; Nakano and Wataru, 2017; Miyata et al., 2018). The mainstay treatment approaches (beta blockers or stellectomy) target the interaction between the ANS and the heart (Wilde et al., 2008; Schwartz and Ackerman, 2022). Moreover, CPVT mice have been shown to have a central nervous system phenotype (Lehnart et al., 2008), and CPVT patients with intellectual disability have a higher risk of arrhythmias (Lieve et al., 2018), indicating that neuronal dysfunction may be associated with a more severe arrhythmic phenotype.

The ANS is a well-established mediator of cardiac function, with the stellate ganglia of the sympathetic chain projecting directly to the myocardium and activation of which leading to increased contractile properties (inotropy, chronotropy). Despite common knowledge of their direct interaction, and the prominent disease feature of adrenergic activity-induced arrhythmias, the stellate ganglia in CPVT have never been investigated before. We hypothesise that there is an unrecognised dysfunction of the ANS which is contributing to the manifestation of the disorder and the fatal arrhythmic phenotype. Our aim in this study was to assess if RyR2 is also expressed in these peripheral, cardiac-modulating (stellate ganglia) neurons, whether RyR2 mutations cause any functional or structural alterations of these neurons, and whether this translates further across the neurocardiac axis to the heart, where it would alter neurotransmitter homeostasis and thus contribute to the manifestation of the disease.

## Methods

### Animals

Wild-type and mutant *Ryr2-R2474S* C57Bl6j mice were purchased from the Jackson Laboratories (031271) and used for the experiments detailed herein - male and female, 2-6 months. Housing, handling, and experiments were performed in agreement with the institutional guidelines.

### Stellate ganglia neuron isolation and culturing

Following terminal anaesthesia (4% isoflurane inhalation in O_2_), stellate ganglia were removed and placed in ice-cold PBS. Single neurons were isolated by an enzymatic dissociation procedure, as previously described (O’Reilly et al., 2024). Ganglia were transferred to a nominally Ca^2+^-free Tyrode’s solution (20°C) (pH 7.4; NaOH) containing (in mM): NaCl 140, KCl 5.4, CaCl_2_ 0.01, MgCl_2_ 1, glucose 5.5, HEPES 5 as well as Liberase TM (26 U/ml) and Elastase (211 U/ml) enzymes. The tissue was gently agitated in a shaking water bath at 37°C for 28 minutes. Subsequently, ganglia were washed in nominally Ca^2+^-free Tyrode’s solution (20°C), and thereafter in normal Tyrode’s solution (20°C) (pH 7.4; NaOH) containing (in mM): NaCl 140, KCl 5.4, CaCl_2_ 1.8, MgCl_2_ 1, glucose 5.5, HEPES 5. Finally, ganglia were transferred to B-27 Plus Neuronal Culture System (Gibco) media (20°C) and single cells were obtained by manual trituration using fire-polished glass pipettes. Single cells were plated on coverslips or 8 well chambers (80807; Ibidi) coated with 100 ug/ml poly-d-lysine and 10 ug/ml laminin. Neurons were left to adhere in a 37°C incubator overnight before experiments were performed. For longer duration culturing, the B-27 Plus Neuronal Culture System media was replenished every 3-4 days, or daily replenishment if performing drug experiments. 2 mice were used for each neuron isolation.

### Cardiac cryosections

Cryosections were prepared according to standard procedures. A Leica CM 1950 cryostat was used to produce 10μm sections from the mid-ventricular region of snap frozen hearts. Cryosections were mounted onto glass slides, left to adhere for 1 hour and then stored at -20°C.

### Immunostaining

After a culturing period, cells were fixed with 4% PFA for 10 min then permeabilized with 0.2% Triton-X in PBS for 10 min. Cryosections were left to thaw for 10 mins, fixed with 4% PFA for 10 min then permeabilized with 0.2% Triton-X in PBS for 60 min. This was followed by a 30 min blocking step in PBS containing 2% glycine, 2% bovine serum albumin (BSA), 0.2% gelatin, and 10% normal goat serum.

The anti-βIII tubulin chicken (AB9354, Sigma-Aldrich, 1:1000), anti-RyR2 rabbit (AB9080, Sigma-Aldrich, 1:200) and anti-tyrosine hydroxylase rabbit (AB152, Sigma-Aldrich, 1:500) primary antibodies were incubated overnight at 4°C. After washing with PBS, the secondary anti-rabbit antibody conjugated with Alexa Fluor 488 (A-11008, Invitrogen, 1:300) and anti-chicken antibody conjugated with Alexa Fluor 568 (A-11041, ThermoFisher, 1:300) were incubated for 1 hour at room temperature. Negative controls were created under the same conditions but without primary antibodies.

**Table 1.** Primary and secondary antibodies used for each experiment/Figure.

| Figure | Primary | Secondary |
| --- | --- | --- |
| 1 | anti-βIII tubulin chicken (1:1000)<br>anti-RyR2 rabbit (1:200) | anti-chicken Alexa Fluor 568 (1:300)<br>anti-rabbit Alexa Fluor 488 (1:300) |
| 3 | anti-βIII tubulin chicken (1:1000) | anti-chicken Alexa Fluor 568 (1:300) |
| 4 | anti-tyrosine hydroxylase rabbit (1:500) | anti-rabbit Alexa Fluor 488 (1:300) |

For detection of RyR2 expression, images were acquired using a Leica Stellaris 5 confocal microscope with a HC PL APO CS2 63x/1.40 oil objective, with 405 (DAPI), 488 (RyR2), and 568 (βIII tubulin) nm laser lines.

For neurite quantification, three pseudo-ganglia areas (regions of interest; ROIs) were randomly selected and imaged. Two rounds of neuron isolation/immunostaining were included in the analysis. Images were acquired using a Leica Stellaris 5 confocal microscope with a HC PL APO CS2 20x/0.75 dry objective, with 405 (DAPI) and 568 (βIII tubulin) nm laser lines. Areas of fluorescence detection were analysed using threshold analysis in ImageJ.

For cardiac innervation quantification, three left ventricular ROIs were imaged per cardiac slice. 3-4 slices were imaged per heart. Four rounds of immunostaining were included in the analysis. Images were acquired using a Leica Stellaris 5 confocal microscope with a HC PL APO CS2 40x/1.30 oil objective, with 488 (TH) nm laser lines. Areas of fluorescence detection were analysed using threshold analysis in ImageJ.

### Calcium homeostasis

Isolated stellate ganglia neurons were cultured overnight then loaded with 10μM Fura Red AM (ThermoFisher, F3021) in B-27 Plus Neuronal Culture System media for 30 mins in a 37°C, 5% CO_2_ incubator. Cells were washed and stored in normal Tyrode’s solution. Live-cell fluorescence imaging was performed using a Leica DMi8 microscope in a 37°C, 5% CO_2_ chamber with a HC PL FLUOTAR 63x/1.30 oil objective and 488 nm laser line. 1 min recordings were measured, with 10mM caffeine (Sigma-Aldrich, C0750; in normal Tyrode’s solution) application to the recording chamber around 10 seconds after the start of recording. Fluorescence intensity over time was analysed using ImageJ.

### Neurotransmitter quantification

Neurotransmitter quantification was performed by Creative Proteomics^TM^.

#### 1. Sample Information

10 selected CPVT mouse heart ventricular tissue samples were quantitatively analyzed for the measurement of epinephrine (EPI), norepinephrine (NE), and normetanephrine (NMN) by using liquid chromatography-mass spectrometric multiple reaction monitoring (LC MRM/MS).

#### 2. Standard Preparation

Calibration solution: a standard solution was prepared in 70% acetonitrile with reference substances of all targeted metabolites. This solution was further diluted step by step to have 10 calibration solutions in 70% acetonitrile. The concentrations of each compound ranged from 0.000001 to 100 μM.

#### 3. Sample Preparation and Analytical Method

Frozen tissues were precisely weighed into 2-mL homogenization tubes. 3 μL of water per mg of tissue and two metal beads were added to each tube. The samples were homogenized on a MM 400 mixer mill for 2 min at 30 Hz. 7 μL of acetonitrile per mg of raw tissue was then added. The samples were homogenized again for 3 min, followed by ultra-sonication in an ice-water bath for 2 min. The samples were centrifuged at 21,000 g and 10 °C for 10 min. The clear supernatants were collected for following LC-MRM/MS. 20 μL of each of the above solutions was mixed with 20 μL of an isotope-labeled internal standard solution, 60 μL of 20-mM dansyl chloride solution and 20 μL of a pH buffer. The mixtures were incubated at 40 °C for 45 min. 10 μL aliquots were injected into a 15-cm C18 column to run UPLCMRM/MS on an Agilent 1290 UHPLC system coupled to an Agilent 6495C QQQ mass spectrometer. 0.1% formic acid in water and in acetonitrile were used as the binary solvents for gradient elution under optimized separation and detection conditions.

#### 4. Analytical Results

In the above assays, concentrations of the detected analytes were calculated with internal standard calibration.

### Statistics

Statistical analysis was carried out with GraphPad Prism version 8.4.3(686) for Windows (GraphPad Software). Data are shown as mean ± SEM. Normality was tested by the Kolmogorov-Smirnov test. An unpaired Student’s t-test was used in the case of normally distributed data. A Mann-Whitney test or a Wilcoxon rank test was applied when normality and/or equal variance test failed. *P*<0.05 was considered statistically significant.

## Results

### RyR2 is abundantly expressed in stellate ganglia

Analysis of published RNA sequencing datasets (Bayles et al., 2018; Sharma et al., 2023) revealed that *Ryr2* is abundantly expressed in mouse stellate ganglia – both when RNA sequencing was performed on isolated stellate ganglia neurons (Figure 1Ai) and whole stellate ganglia (Figure 1Aii). We found *Ryr2* to be more abundantly expressed than the characteristic neuronal markers *Chat, Syn1* and *Scn8a* (Figure 1A).

**Figure 1.**
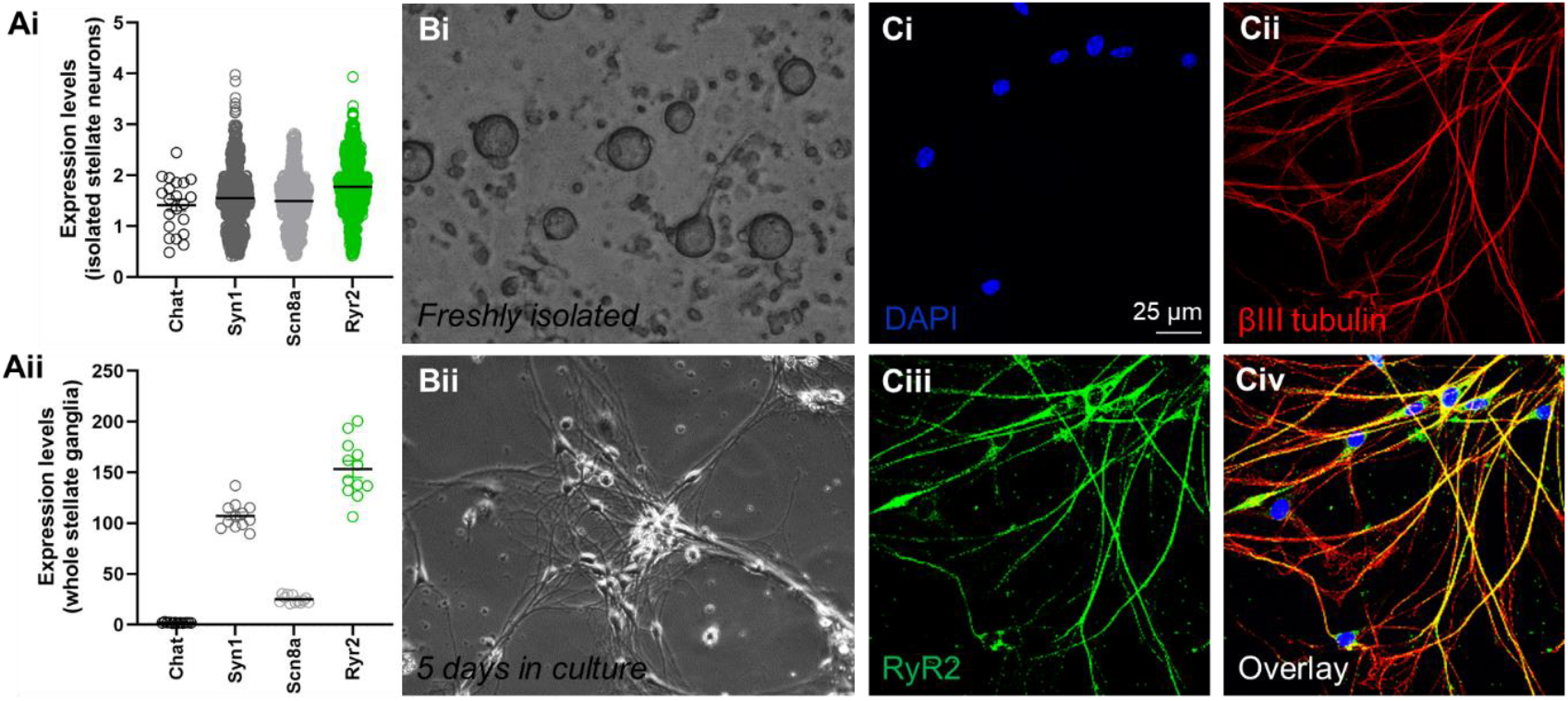
RyR2 is abundantly expressed in mouse stellate ganglia neurons. A) Published RNA sequencing datasets showing *Ryr2* expression levels in isolated mouse stellate ganglia neurons (Ai) and whole mouse stellate ganglia (Aii). Expression levels of *Chat, Syn1* and *Scn8a* are shown for comparison. B) Light microscopy showing the appearance of freshly isolated stellate ganglia neurons (Bi) and those neurons cultured for 5 days (Bii). C) Confocal microscopy of cultured stellate ganglia neurons stained for DAPI (Ci, blue), βIII tubulin (Cii, red) and RyR2 (Ciii, green). Civ) Shows the overlay. Scale bar is 25 μm.

Using immunocytochemistry on isolated wildtype (WT) mouse stellate ganglia neurons, we revealed for the first time that RyR2 protein is expressed in these peripheral, cardiac-modulating neurons. Figure 1B shows the appearance of these neurons when freshly isolated (Figure 1Bi) and after 5 days in culture (Figure 1Bii) – a time duration which enables neurons to regrow their axons and dendrites. Immunostaining of cultured neurons with the neuronal marker βIII tubulin (Figure 1Cii) revealed that RyR2 protein was detectable throughout the neuron structure, from cell bodies through to dendritic arborisations (Figure 1Ciii-iv).

### The CPVT *Ryr2*-R2474S mutation leads to functional alterations of stellate ganglia neuron calcium homeostasis

Having established the presence of RyR2 in stellate ganglia neurons, we next investigated its functional relevance in the setting of a CPVT mutation employing *Ryr2*-R2474S mice. In cardiomyocytes of these mice, the mutation is known to cause calcium leakage (Lehnart et al., 2008). Functional investigation of calcium homeostasis revealed a dysregulation of calcium handling in isolated CPVT stellate ganglia neurons. Isolated neurons were loaded with the fluorescent calcium indicator Fura Red AM and exposed to 10mM caffeine to release the calcium stored in the intracellular stores (the endoplasmic reticulum). Representative fluorescence images are shown in Figure 2A and F/F0 traces in Figure 2B. The maximal reduction of fluorescence intensity after caffeine exposure - a measure of total intracellular calcium release - was found to be significantly decreased in *Ryr2*-R2474S neurons (F/F0 WT 0.45±0.01 vs CPVT 0.53±0.01, p<0.01) (Figure 2C). Hence, there is a functional effect of the *Ryr2-* R2474S mutation in CPVT stellate ganglion neurons, with potential implications for calcium leakage.

**Figure 2.**
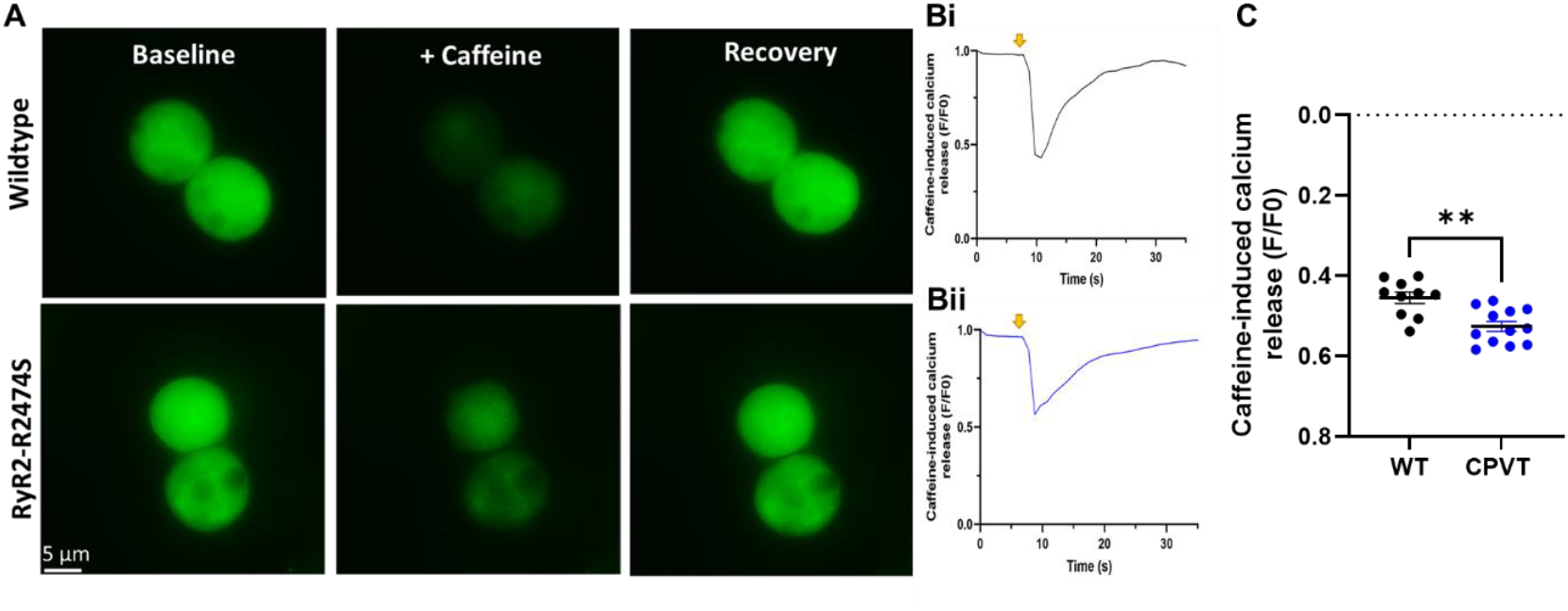
*Ryr2-R2474S* stellate ganglia neurons have altered calcium handling. A) Live cell widefield fluorescence microscopy of isolated stellate ganglia neurons cultured overnight, loaded with the intracellular calcium indicator Fura Red AM (10μM), and exposed to 10mM caffeine. B) Representative F/F0 traces showing fluorescence changes over time upon caffeine exposure in wildtype (Bi) and *Ryr2*-R2474S neurons (Bii). Arrows indicate the addition of caffeine. C) Quantification of the maximal reduction of fluorescence intensity revealed a significantly reduced reduction in *Ryr2-R2474S* neurons (**p<0.01) (unpaired t-test). Scale bar is 5 μm.

### Alterations in neurite outgrowth from *Ryr2*-R2474S stellate ganglia neurons

Calcium handling in neurons is known to influence dendritic development (Konur and Ghosh, 2005), so we next quantified neurite outgrowth from cultured CPVT stellate ganglia neurons. Neurons were isolated and cultured for 7 days before being stained for DAPI (blue) and βIII tubulin (red) (Figure 3A). The DAPI-positive area was similar (WT 12.4±3.6 vs CPVT 11.1±0.9), indicating similar overall cell densities (Figure 3B). The βIII tubulin-positive area was significantly increased in *Ryr2*-R2474S cultures (WT 43.4±6.0 vs CPVT 75.2±5.6, p<0.01) (Figure 3C), and remained significantly increased after normalising to DAPI-positive area (WT 4.1±0.6 vs CPVT 6.9±0.6, p<0.01) (Figure 3D), indicating that cultured *Ryr2*-R2474S neurons showed increased neurite outgrowth. To provide further evidence for a role of RyR2 in this process, we performed the same measurements following pharmacological induction of RyR2 leakage, thereby mimicking the CPVT condition. Incubation of WT cultured neurons with 500nM ryanodine for 7 days caused a significant increase in neurite outgrowth when compared to vehicle (H_2_O) control (vehicle 4.3±0.3 vs ryanodine 5.5±0.3, p<0.05) (Figure 3E).

**Figure 3.**
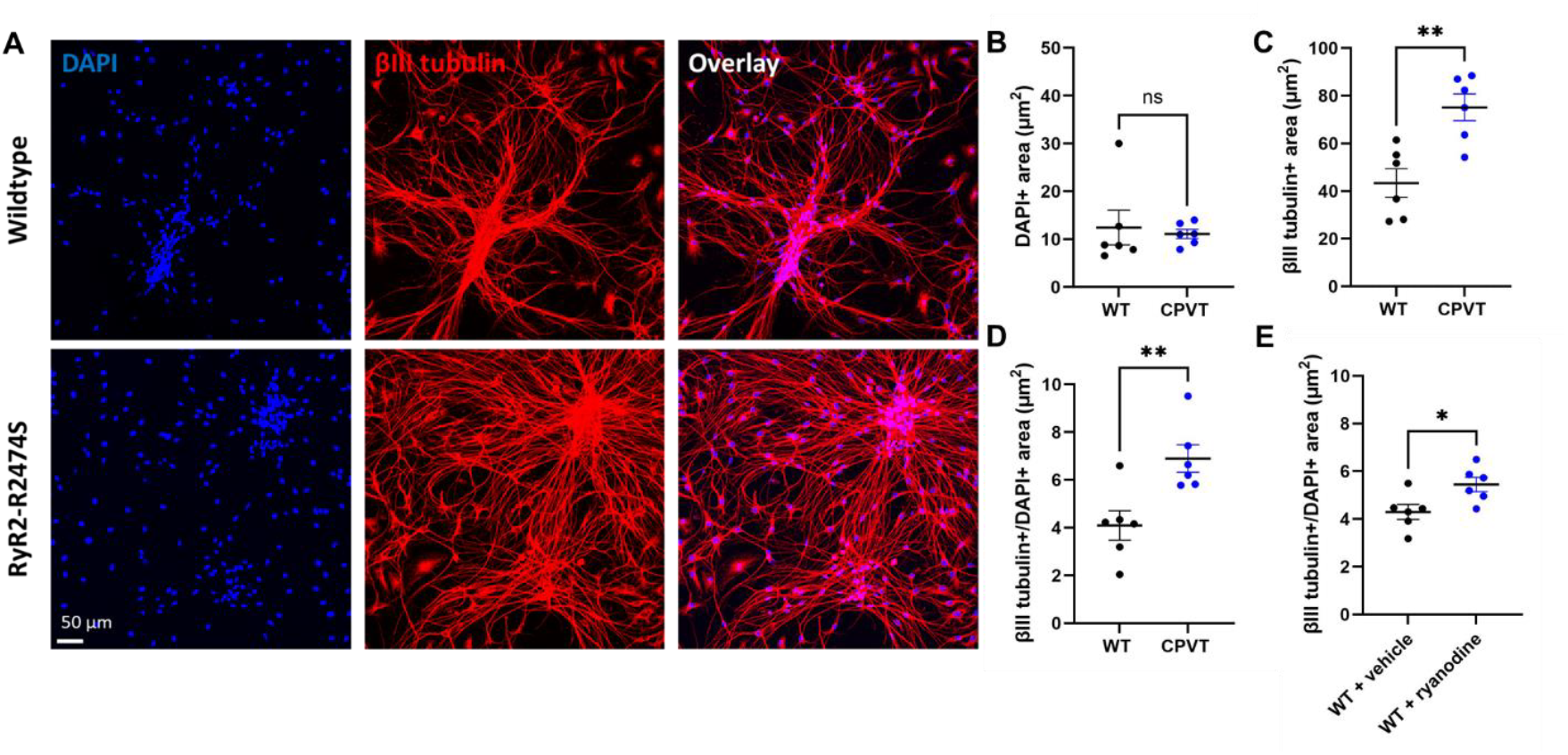
Cultured *Ryr2*-R2474S stellate ganglia neurons have increased neurite outgrowth. A) Confocal imaging of cultured stellate ganglia neurons stained for DAPI (blue) and βIII tubulin (red). B) The DAPI-positive area was similar, indicating similar cell densities. C) The βIII tubulin-positive area was significantly increased in *Ryr2*-R2474S cultures (**p<0.01). D) Quantification of βIII tubulin-positive area normalised to DAPI-positive area revealed a significant increase of neurite outgrowth from *Ryr2*-R2474S cell bodies compared to wildtype (WT) (**p<0.01) (C). Quantification of βIII tubulin-positive area normalised to DAPI-positive area in WT neurons cultured for 7 days with either 500nM ryanodine or vehicle H_2_O control revealed a significant increase of neurite outgrowth with ryanodine exposure (*p<0.05) (unpaired t-tests). Scale bar is 50 μm.

### Sympathetic hyperinnervation in ventricular myocardium of *Ryr2*-R2474S mice

We next investigated whether the observed increase in neurite outgrowth was associated with altered sympathetic innervation in CPVT mouse hearts. Neuronal innervation density was assessed by staining 10 μm cryosections of mid-ventricular myocardium for the sympathetic neuron marker tyrosine hydroxylase (TH) (Figure 4A). Quantification of the TH-positive area revealed a significant increase in the density of sympathetic neuron innervation in ventricular tissue of *Ryr2*-R2474S hearts (Figure 4B-D). Hyperinnervation was consistently observed throughout the left ventricular sub-epicardium area, as evidenced by statistical comparison of individual regions of interest (ROIs) (WT 2.0±0.2 vs CPVT 4.0±0.3, p<0.0001, Figure 4B), individual slices (WT 1.7±0.2 vs 3.8±0.3, p<0.0001, Figure 4C) and when performing whole heart analysis (WT 1.7±0.3 vs CPVT 3.9±0.5, p<0.01, Figure 4D).

**Figure 4.**
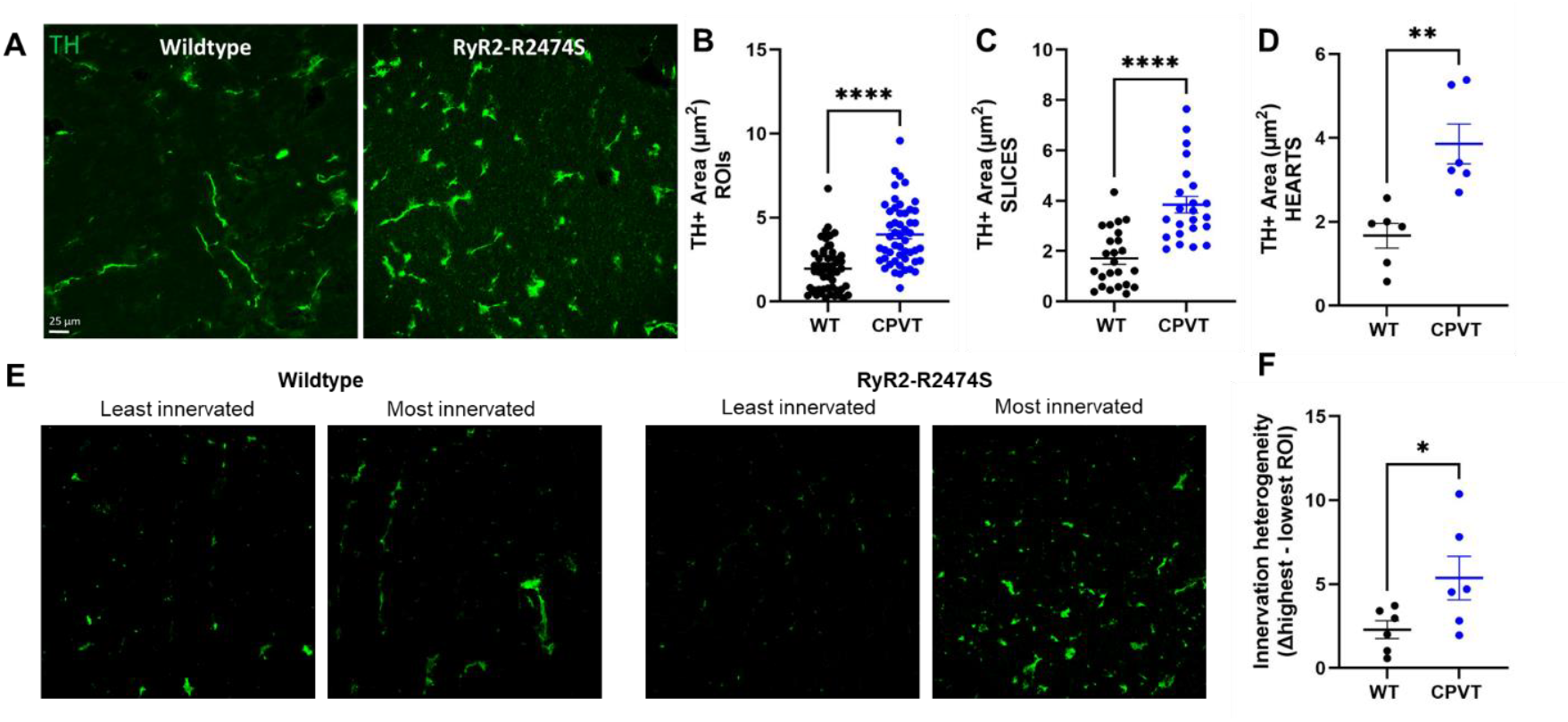
*Ryr2-R2474S* ventricular myocardium is sympathetically hyper-innervated. A,E) Confocal imaging of mid-ventricular heart cryosections stained for the sympathetic neuron marker tyrosine hydroxylase (TH). B-D) Quantification of the TH-positive area revealed a significant increase of sympathetic neuron density in individual *Ryr2*-R2474S regions of interest (ROIs) (****p<0.0001) (B), individual slices (****p<0.0001) (C) and whole hearts (**p<0.01) (D). An increased heterogeneity of cardiac innervation was also evident from comparison of the most hypo- and hyper-innervated areas of each heart (*p<0.01) (F) (unpaired t-tests). Scale bar is 25 μm.

Moreover, there was a significant increase in the heterogeneity of this augmented innervation, with greater variability in *Ryr2*-R2474S hearts as evidenced by comparison of the most hypo- and hyper-innervated regions of interest (2.3±0.5 vs 5.4±1.3, p<0.05) (Figure 4E-F).

### The *Ryr2*-R2474S mutation leads to reduced epinephrine levels and increased norepinephrine turnover in ventricular myocardium

Neuronal innervation is the major determinant of neurotransmitter release within the heart, which is the ultimate functional link between the ANS and the cardiac system. To investigate any alterations in the setting of CPVT, we performed neurotransmitter quantification with LC MRM/MS. Quantification of the concentration of neurotransmitters and their metabolites in ventricular tissue revealed that in *Ryr2*-R2474S hearts there are reduced levels of epinephrine (WT 0.041±0.005 vs 0.023±0.002, p<0.01), similar levels of norepinephrine (NE), an elevated level of the NE metabolite normetanephrine (NMN), and consequently a significant increase in the NMN/NE ratio – a known index of NE turnover (WT 0.166±0.007 vs CPVT 0.268±0.012, p<0.01) (Figure 5). This suggests that the observed sympathetic hyperinnervation in CPVT leads to increased neurotransmitter turnover within the heart.

**Figure 5.**
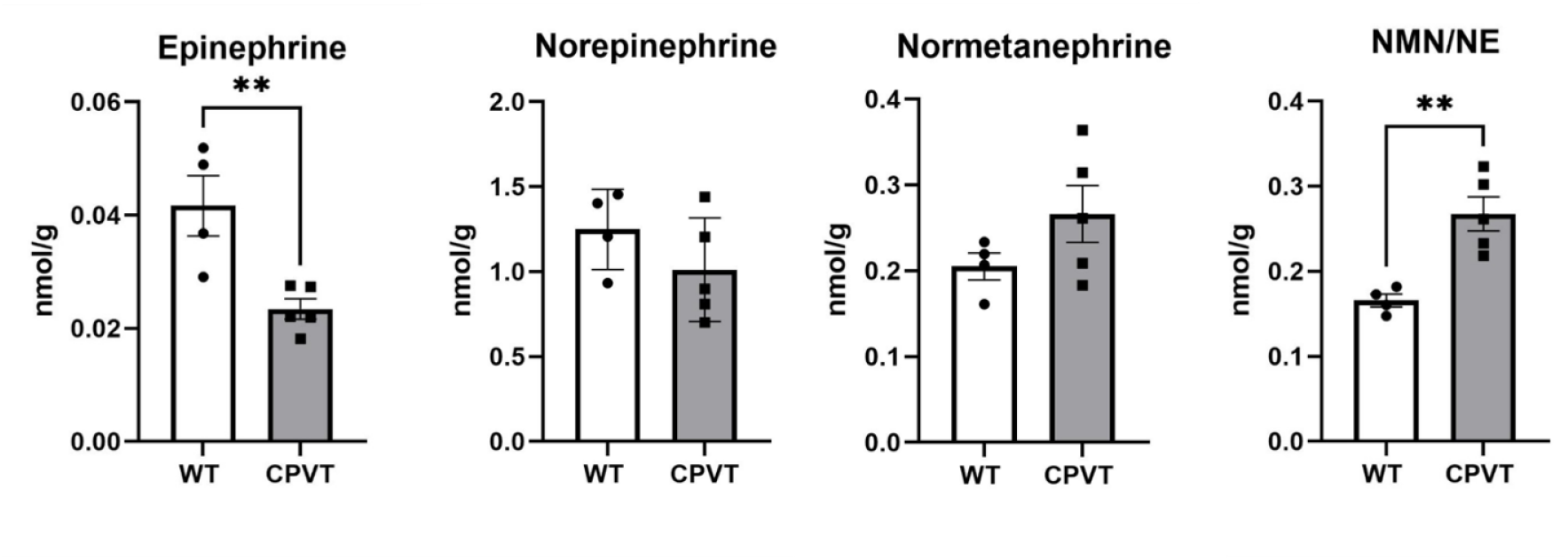
*Ryr2*-R2474S ventricular tissue has reduced levels of epinephrine and elevated levels of norepinephrine turnover. Neurotransmitter quantification using LC MRM/MS showed that in *Ryr2*-R2474S ventricular tissue there is reduced levels of epinephrine (**p<0.01), similar levels of norepinephrine (NE), elevated levels of normetanephrine (NMN) and a significant increase in the NMN/NE ratio, indicating increased NE turnover in *Ryr2*-R2474S hearts (*p<0.01) (unpaired t-tests).

## Discussion

Here we show for the first time that RyR2, previously thought to only be present in heart and brain tissue, is also very abundantly expressed in peripheral, cardiac modulating neurons (those of the stellate ganglia of the sympathetic chain). We further reveal that *RyR2* mutations that are known to cause CPVT (R2474S) lead to functional alterations of these stellate ganglia neurons. The *Ryr2*-R2474S mutation leads to reduced caffeine-induced calcium release, which is indicative of there being less calcium stored in intracellular stores and strongly suggests the presence of calcium leakage, similarly to in cardiomyocytes. Consistent with the known effects of neuronal calcium leakage (Konur and Ghosh, 2005), we found that these neurons also have structural alterations in the way of increased neurite outgrowth when maintained in culture. Not only did we observe functional and structural alterations of the sympathetic nervous system, we furthermore revealed that these changes extended towards the heart, where sympathetic hyper-innervation of the CPVT ventricular myocardium was observed. Lastly, we found that these broad autonomic alterations culminated in crucial changes in the concentrations of neurotransmitters and their metabolites in ventricular tissue - specifically changes in (nor)epinephrine, their metabolites and respective ratios, which have a fundamental role in regulating cardiac activity (Lepeschkin et al., 1960; Kaumann et al., 1989).

Until now, arrhythmia research in CPVT has focussed predominantly on cardiomyocytes. The prevailing dogma of CPVT is that mutant RyR2 on the sarcoplasmic reticulum membrane spontaneously leaks calcium into the cytoplasmic space due to various underlying mechanisms, which then triggers arrhythmias (Priori et al., 2021). This aspect of the arrhythmic hypothesis is not disputed. However, what we add here is an additional mechanism underlying the increased propensity for RyR2 calcium leakage. RyR2 releases calcium when a threshold of calcium inside the SR is reached. We propose that in CPVT, the sympathetic hyper-innervation and increased NE turnover magnifies SR calcium overload during sympathetic activation, thereby facilitating spontaneous calcium release and pro-arrhythmia. Hence, the neuronal aspect of the condition contributes significantly to the arrhythmic phenotype, which is why treatments targeted towards the excess NE beta receptor activation (beta blockers) and sympathetic hyper-innervation (stellectomy) are the most effective treatment approaches. *RYR2* CPVT may therefore be redefined as a neurocardiac condition.

A recent study revealed altered neuronal innervation in the hearts of mice with arrhythmogenic cardiomyopathy (Vanaja et al., 2024). However, in structurally abnormal hearts, it is unclear what the progression of the disease is - whether the neuronal and cardiac dysfunction occurs simultaneously, or whether cardiac structural remodelling drives the neuronal changes, in a reverse feedback aspect of the neurocardiac axis. Our current observations in CPVT provide the first evidence of neuronal remodelling in a cardiac channelopathy condition in which the heart is structurally normal. However, it remains possible that dysfunctional cardiomyocytes drive – or at least contribute to – the innervation differences. It is known that cardiomyocytes release neurotrophic factors such as nerve growth factor (NGF) on to innervating neurons across the neurocardiac junction to maintain neuronal innervation (Dokshokova et al., 2022). Hence, altered levels of NGF may (partly) underlie these innervation changes – a speculation which remains to be investigated. It also remains to be addressed whether this redefinition of CPVT as a neurocardiac condition is also applicable to CPVT cases caused by *CASQ2* mutations (the second most commonly associated gene). A similar investigation needs to be performed for confirmation of the conservation of the proposed mechanism.

The results of this investigation have significant clinical potential for both risk stratification and novel treatment design. In ventricular myocardial tissue of *Ryr2-R2474S* mice, we observed an increased normetanephrine to norepinephrine ratio, indicative of enhanced norepinephrine turnover. Hence, the norepinephrine metabolite normetanephrine could be used as a novel blood biomarker (Oeltmann et al., 2004) for assessing the severity of the condition and the likelihood of arrhythmia occurrence – with higher plasma levels indicating augmented cardiac innervation and thus a greater propensity for arrhythmias. A clinical follow-up study could confirm the applicability of these basic science results by comparing measures of autonomic (dys)function in symptomatic vs asymptomatic and mildly versus severely affected patients, for example. This could be achieved with non-invasive measures such as heart rate variability, iodine meta-iodobenzylguanidine (MIBG) imaging, or blood biomarker testing. Moreover, RyR2 stabilisation in the stellate ganglia could pose a novel therapeutic approach. Stabilizing neuronal calcium leakage would reduce neurite outgrowth and cardiac hyperinnervation and thus prevent excess NE from triggering cardiomyocyte calcium leakage.

## Conclusion

The stellate ganglia of the peripheral autonomic nervous system abundantly express RyR2. *RyR2* mutations lead to functional alterations of stellate ganglia neuron calcium handling in the form of calcium leakage and also caused structural alterations in the form of increased neurite outgrowth. This extends to the heart where it induces sympathetic neuron hyperinnervation of the ventricular myocardium. Importantly, this leads to increased cardiac norepinephrine turnover. CPVT may thus be redefined as a neurocardiac disorder, with neuronal dysfunction being a prominent arrhythmia-provoking component of the disease. As such, our findings identify novel avenues for improved disease detection, risk stratification and treatment design.

## Author contributions

Conceptualization, M.O. and C.A.R.; Data collection, M.O.; Data analysis, M.O., A.O.V.; Funding acquisition, M.O. and C.A.R.; Visualization, M.O., A.O.V.; Writing, M.O., A.O.V. and C.A.R.

All authors have read and agreed to the published version of the manuscript.

